# MungeSumstats: A Bioconductor package for the standardisation and quality control of many GWAS summary statistics

**DOI:** 10.1101/2021.06.21.449239

**Authors:** Alan E Murphy, Nathan G Skene

**Affiliations:** UK Dementia Research Institute at Imperial College London

## Abstract

**Summary:** Genome-wide association studies (GWAS) summary statistics have popularised and accelerated genetic research. However, a lack of standardisation of the file formats used has proven problematic when running secondary analysis tools or performing meta-analysis studies. To address this issue, we have developed MungeSumstats, a Bioconductor R package for the standardisation and quality control of GWAS summary statistics. MungeSumstats can handle the most common summary statistic formats, including variant call format (VCF) producing a reformatted, standardised, tabular summary statistic file, VCF or R native data object.

**Contact:** Alan Murphy, Nathan Skene

**Availability and implementation:** MungeSumstats is available on Bioconductor (v 3.13) and can also be found on Github at: https://neurogenomics.github.io/MungeSumstats

**Supplementary information:** The analysis deriving the most common summary statistic formats is available at: https://al-murphy.github.io/SumstatFormats

## Introduction

Genome-wide association studies (GWAS) summary statistics are used to distribute the most important outputs of GWASs in a manner which does not require the transfer of individual-level personally identifiable information from participants. Summary statistics from past studies tend to become more valuable over time as it becomes possible to meta-analyse and integrate them with new annotation information through approaches such as Linkage Disequilibrium Score Regression (LDSC)(1), Generalized Gene-Set Analysis of GWAS Data, MAGMA(2), and multi-phenotype investigations(3,4). Summary statistics are also commonly integrated for use in the meta-analysis of GWAS. However, these tools and this integration require a standardised data format which was historically lacking from the field. The diversity of data formats in summary statistics has been a result of the phenotypes in question, for example disease-control or quantitative trait, the software used to perform the analysis, such as PLINK(5) and GCTA(6) or just the preference of the consortium in question.

There have been movements to standardise the summary statistic file format such as the NHGRI-EBI GWAS Catalogue standardised format(7) and the SMR Tool binary format(8). More recently, the variant call format to store GWAS summary statistics (GWAS-VCF)(9) has been developed which has manually converted over 10,000 GWAS to this format. While GWAS-VCF offers a standardised format that future GWAS consortium may adopt, there are still a multitude of past, publicly available GWAS which have not been standardised(10)(11)(12)(13). For instance, although their summary statistics are publicly available, the GWAS for Cerebral small vessel disease(14) is not yet available in VCF format via IEU GWAS. Furthermore, as VCF is not yet the standard for sharing files between geneticists, unpublished GWAS shared internally within genetics consortia or provided by personal genetics companies are still found in a variety of summary statistic formats. As such, there is a need for tools to move between the various formats in which summary statistics are stored.

The standardisation of GWAS summary statistics also requires quality control to ensure cohesive integration. For example, checking if the non-effect allele from the summary statistics matches the reference sequence from a reference genome to ensure consistent directionality of allelic effects across GWAS. In addition, downstream analysis tools often require a degree of quality control which, in the case of meta-analysis, must be applied across all GWAS. One such example is the removal of all non-biallelic SNPs is a common requirement of all downstream analysis(9).

To address these issues, we introduce MungeSumstats a Bioconductor R package for the rapid standardisation and quality control of many GWAS summary statistics. MungeSumstats can handle the most common summary statistic formats as well as GWAS-VCFs to enable the integrative meta-analysis of diverse GWAS. MungeSumstats also offers a comprehensive and tuneable quality control protocol with defaults for common, best-practice approaches. MungeSumstats capitalises on R’s familiar interface, is readily accessible through Bioconductor and utilises an intuitive approach, running with a single line of input code.

## Heterogeneity in GWAS Formats

To demonstrate the diversity in summary statistics across GWAS, we analysed a public repository of over 200 publicly available GWAS(15). From this, the most common summary statistics were derived, see Figure 1 for the 12 most common file header formats.

**Figure 1:**
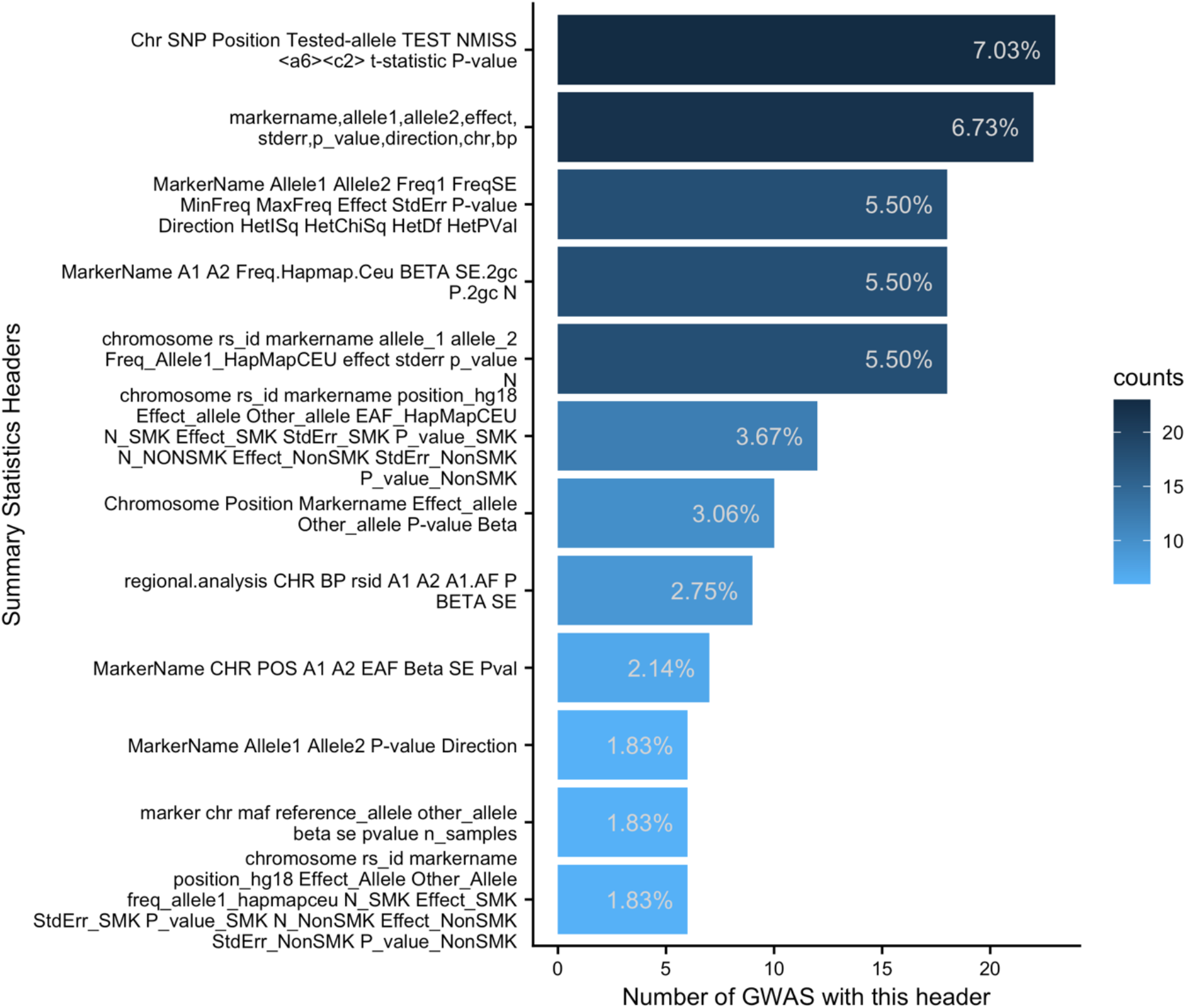
Most common summary statistic formats. shows the most common summary statistic formats from a repository of over 200 publicly available GWAS(15). Note that a GWAS can have more than 1 summary statistics file and ‘<a6><c2>‘ is the symbol ‘¶¬‘ read into R.

A total of 327 summary statistic files were derived from the analysis which corresponded to 127 unique formats. Thus on average, every 2.5 summary statistic files had a unique format, showing the clear disparity across GWAS. The 12 most common formats, shown in Figure 1, accounted for approximately 47% all summary statistics.MungeSumstats has been tested on these 12 most common formats and is able to standardise their summary statistics.

## Implementation

MungeSumstats was implemented using the R programming language (v 4.0) and Bioconductor S4 data infrastructure (v 3.13) enabling the full analysis of summary statistics within the R environment. The package removes the need for external software to perform the standardisation and quality control steps.

MungeSumstats’ implementation ensures both memory and speed efficiency through the use of R data.table (v.1.14.0)(16), which can take advantage of multi-core parallelization. Moreover, MungeSumstats benefits from Bioconductor’s infrastructure for efficient representation of full genomes and their SNPs, using BSgenome (v 1.59.2) SNP reference genomes(17). Either Ensembl’s GRCh37 or GRCh38 are queried dependent on the build for the particular GWAS. Numerous of MungeSumstats’ quality control steps for summary statistics require the use of a reference genome. For example, an allele flipping test is run, see Table 1, to ensure consistent directionality of allelic effect and frequency variables. The effect or alternative allele is always assumed to be the second allele (A2), in line with the approach for GWAS-VCF(9). Moreover, MungeSumstats can impute any missing, essential information like SNP ID, base-pair position and effect/non-effect allele.

**Table 1:** MungeSumstats Implemented Checks. lists the quality control and standardisation checks conducted. Most checks are optional and can be set by the user. Here CHR is chromosome, BP is Basepair position, A1 is the non-effect allele, A2 is the effect allele, N is the sample size, INFO is imputation information score, FRQ is the minor allele frequency (MAF) of the SNP, SNP ID is the single nucleotide polymorphism reference ID, P is the unadjusted p-value, Z is z-score, OR is odds ratio, LOG_ODDS is the log odds ratio, BETA is the effect size estimate relative to the alternative allele and SIGNED_SUMSTAT is the directional effect size estimate for the summary statistics.

| Step | MungeSumstats Check | Description |
| --- | --- | --- |
| 1 | Check VCF format | If the input file is in variant call format (VCF), if so import |
| 2 | Check tab, space or comma delimited | If input is space or comma delimited convert to tab delimited. Can handle .tsv, .txt, .csv, .tsv.gz, .txt.gz, .csv.gz, .tsv.bgz, .txt.bgz, .csv.bgz, .vcf, .vcf.gz, .vcf.bgz files. |
| 3 | Check for header name synonyms | If any alternative names are found for SNP, BP, CHR, A1, A2, P, Z, OR, BETA, LOG_ODDS, SIGNED_SUMSTAT, N, N_CAS, N_CON, NSTUDY, INFO or FRQ convert them to a standard name. Robust conversion approach with 176 unique mappings |
| 4 | Check for multiple models or traits in GWAS | If multiple, user must specify one to analyse |
| 5 | Check for uniformity in SNP ID | Ensure no mix of RS ID, missing 'rs' prefix and/or CHR:BP |
| 6 | Check for CHR:BP:A2:A1 all in one column | Split into separate columns if found |
| 7 | Check for CHR:BP in one column | Split into separate columns if found |
| 8 | Check for A1/A2 in one column | Split into separate columns if found |
| 9 | Check if CHR and/or BP is missing | If so, infer from the chosen reference genome |
| 10 | Check if SNP ID is missing | If so, infer from the chosen reference genome |
| 11 | Check if A1 and/or A2 are missing | If so, infer from the chosen reference genome |
| 12 | Check that vital columns are present | Check for the necessary columns; SNP, CHR, BP, P, A1, A2 |
| 13 | Check for one signed/effect column | Effect columns Z, OR, BETA, LOG_ODDS, SIGNED_SUMSTAT |
| 14 | Check for missing data | If data is missing from any entry, remove the SNP |
| 15 | Check for duplicated columns | If there are any remove one |
| 16 | Check for p-values lower than 5e-324 | These are not recognised in R and cause issues with downstream analysis software like LDSC/MAGMA. User can convert to 0. |
| 17 | Check N column | Ensure it is an integer and check if the sample size for a SNP isn't greater than mean multiplied by five times the standard deviation. Removes SNPs that have substantial more samples than the rest. |
| 18 | Check SNPs are RS ID's | Checks validity of SNP IDs as RS IDs, other IDs can still be used |
| 19 | Check for duplicated rows, based on SNP ID | Duplicates are removed |
| 21 | Check for duplicated rows, based on base-pair position | Duplicates are removed |
| 22 | Check for SNPs on reference genome | Correct any missing from reference genome using BP and CHR |
| 23 | Check INFO score | Remove SNPs with imputation score less than 0.9 |
| 24 | Check for strand-ambiguous SNPs | Remove strand-ambiguous SNPs if found |
| 25 | Check for non-biallelic SNPs (infer from reference genome) | Infer from chosen reference genome and remove any if found |
| 26 | Check for allele flipping | The effect/alternative/minor allele is assumed to be A2. The allele flipping function checks A1 against a reference genome. For a given SNP, if A1 doesn't match the reference genome sequence (i.e. it is the alternative allele, not the reference allele for example), A1 and A2 along with the effect and frequency columns are flipped, creating consistent directionality of allelic effects across GWAS. |
| 27 | Check for SNPs on chromosome X, Y, and mitochondrial SNPs (MT) | If any are found these are removed. |
| 28 | Check output format is LDSC ready | Standardised file can be passed to LDSC without pre-processing |
| 29 | Check effect column values | Ensure effect columns (like BETA) aren't equal to 0 |
| 30 | Check Standard Error | Ensure standard error (SE) is positive |
| 31 | Check dropped and imputed values | Return indicators of the imputed values for a SNP and return the SNPs and the reason for exclusion because of QC. |

Using these two infrastructures, MungeSumstats conducts more than 30 checks on the inputted summary statistics file, see Table 1 for a description of their use. MungeSumstats is also written to ensure the ease of addition of further checks so if users have summary statistics which can’t currently be handled in MungeSumstats, these can be incorporated easily in future releases. Finally, MungeSumstats returns a reformatted, tabular summary statistics file, a VCF or an R native data object (data.table, VRanges or GRanges) with standardised columns for the information necessary for downstream analysis.

## Usage

Once MungeSumstats is installed, usage involves a single line of code or one function call (*format sumstats*) with the path to the summary statistics file of interest. Then, the path to the reformatted, standardised summary statistic file is returned. MungeSumstats also offers adjustable parameters to manage the quality control steps. These include options to adjust the imputation information score (INFO) cut-off threshold, the number of samples (N) outliers cutoff threshold and whether to remove mitochondrial SNPs or SNPs on the X or Y chromosome (see Table 1). Quality control steps which use a reference genome can also be adjusted such as whether to filter SNPs based on their RS ID’s presence on the reference genome, whether to check for allele flipping and whether to remove multi-allelic or strand-ambiguous SNPs. These parameters ensure MungeSumstats can be adjusted to the user’s analysis pipelines.

## Conclusion

Here, we presented MungeSumstats, a Bioconductor package for the standardisation and quality control of GWAS summary statistics. This package enables integration of summary statistics of vastly different formats, simplifying meta-analysis and summary statistics use in other secondary research applications. The package provides an efficient, user-friendly R-native approach, returning a standardised, tabular format file, VCF or R native data object. This ensures that the summary statistics are accessible to the average user. Moreover, MungeSumstats is written to permit future development of additional standardisation steps if users encounter issues with their specific GWAS.

## Acknowledgements

This work was supported by a UKDRI Future Leaders Fellowship [grant number MR/T04327X/1] and the UK Dementia Research Institute which receives its funding from UK DRI Ltd, funded by the UK Medical Research Council, Alzheimer’s Society and Alzheimer’s Research UK. We thank Alexandru Voda (@alexandruioanvoda) for contributing via github in making an early version of this package work across operating systems.

## Data availability

The data to derive the summary statistic formats is open source and collated at: https://github.com/mikegloudemans/gwas-download

